# Identification of key regulators in Prostate cancer from gene expression datasets of patients

**DOI:** 10.1101/643643

**Authors:** Irengbam Rocky Mangangcha, Md. Zubbair Malik, Ömer Küçük, Shakir Ali, R.K. Brojen Singh

## Abstract

Identification of key regulators and regulatory pathways is an important step in the discovery of genes involved in cancer. Here, we propose a method to identify key regulators in prostate cancer (PCa) from a network constructed from gene expression datasets of PCa patients. Overexpressed genes were identified using BioXpress, having a mutational status according to COSMIC, followed by the construction of PCa Interactome network using the curated genes. The topological parameters of the network exhibited power law nature indicating hierarchical scale-free properties and five levels of organization. Highest degree *hubs* (*k*≥65) were selected from the PCa network, traced, and 19 of them were identified as novel key regulators, as they participated at all network levels serving as backbone. Of the 19 hubs, some have been reported in literature to be associated with PCa and other cancers. Based on participation coefficient values most of these are *connector* or *kinless hubs* suggesting significant roles in modular linkage. The observation of non-monotonicity in the rich club formation suggested the importance of intermediate hubs in network integration, and they may play crucial roles in network stabilization. The network was self-organized as evident from fractal nature in topological parameters of it and lacked a central control mechanism.

## Introduction

Prostate is a gland of the male reproductive system which secretes seminal fluid in human adult (1). According to World Health Report 2014, the cancer of prostate or Prostate cancer (PCa) in man is second most common cancer, after lung cancer, and is responsible for a fifth of cancer deaths in males worldwide (2). PCa, based on the type of origin in prostate, can be classified into five types: (i) acinar adenocarcinoma, (ii) ductal adenocarcinoma, (iii) transitional cell (or urothelial) cancer, (iv) squamous cell cancer and (v) small cell prostate cancer, with adenocarcinomas being the most common, even though metastasis is much quicker in other types (3, 4).

In recent years, gene expression studies using high-throughput techniques namely next generation sequencing, microarray and proteomics have led to the identification of new genes and pathways in PCa. The identification of novel key regulators is important as the current therapeutic modalities against PCa, including the use of antiandrogens and blocking androgen synthetic pathway (5) and using Luteinizing hormone-releasing hormone (LHRH) agonists and antagonists along with cytotoxic anticancer drugs, cause notable side effects (6,7). Moreover, PCa diagnosis, which is largely dependent on the Prostate specific antigen (PSA) and Digital rectal examination (DRE), has its own limitations (8,9). PSA is also elevated in benign prostatic hyperplasia and other noncancerous conditions (8). This necessitates the discovery of more reliable biomarkers for better and early diagnosis, as well as identification of new targets other than the genes involved in androgen metabolism for the discovery and development of new and more potent drugs which have less toxicity and lesser side effects.

Genes are regulated in a coordinated way and the expression of one gene usually depends on the presence or absence of another gene (gene interaction). Network theory, which studies the relations between discrete objects through graphs as their representations, can be used to study complex gene regulatory networks which can have different types (random, scale-free, small world and hierarchical networks). The development of algorithms to study of these networks can provide an important tool to find/identify disease-associated genes in complex diseases such as cancer. Earlier, the network theory-based methods have been used to predict disease genes from networks generated using curated list of genes reported to be associated with the disease and mapping them to the human gene interaction network (HPRD database) (10). In such approach, the studies have been limited to the curated gene list forming the network not completely representing the system and patient-specific information is not considered. Moreover, current studies on complex networks in human disease models to discover key disease genes rely mostly on clustering and identifying the high degree hubs or/and motif discovery from the networks (11,12). Therefore, the application of network theoretical methods to the PPI networks of cancer associated genes constructed from the corresponding genes by analyzing high-throughput gene expression datasets of human cancer patients may be used for better sensitivity and forecast in understanding the key regulating genes of the corresponding disease. The clinical impact of using patients’ gene expression data over gene expression data from cancer cell lines will also give a systematic insight in predicting key regulator genes expressed in cancer and understanding their roles in disease manifestation and progression. In this study, we have used the gene expression data (RNAseq) of PCa patients to construct complex PPI network and analyze it. The study gives equal importance to the *hubs*, *motifs* and *modules* of the network to identify the key regulators and regulatory pathways not restricting only to overrepresented *motifs* or *hubs* identification, establishing a relationship between them in gene-disease association studies using network theory. The method used in this study is new and takes a holistic approach for predicting key disease genes and their pathways within network theoretical framework using datasets of PCa patients.

## Materials and methods

### Identification and selection of PCa-associated genes

BioXpress v3.0 (https://hive.biochemistry.gwu.edu/bioxpress), which uses TCGA (https://tcga-data.nci.nih.gov/) RNA sequencing datasets derived from the human cancer patients (13,14), was used to differentiate the deregulated genes in cancer. The cancer browser tool of COSMIC (https://cancer.sanger.ac.uk/cosmic) (15) was used for the mutational status and accordingly, non-redundant genes overexpressed in human PCa were identified.

### Construction of Protein-protein interaction (PPI) network

After excluding the redundancy and redundant copies, out of 4,890 genes found to be significantly overexpressed (***FC*** > 1, adjusted ***p*** < 0.05) in PCa patients from BioXpress, 3,871 genes, which had mutational status in PCa according to COSMIC, were used to construct an interactome network using GeneMania app (16) in Cytoscape 3.6.0 (17). From the network, only the physical interaction network, which represented the Protein-protein network of PCa-associated genes, was extracted. After curation of the network (removal of isolated node/nodes), a PPI network of 2,960 nodes and 20,372 edges was finally constructed as Primary network representing a graph denoted by *G(N,E)*, where, *N* is the set of nodes with *N={n_i_}; i=1,2,.,N* and *E* the set of edges with *E={e_ij_}; i,j=1,2,3,….,N*.

### Method for detection of levels of organization

Considering the size of the network and its sensitivity, Louvain method of modularity (***Q***) maximization was used for community detection (18). The first level of organization was established by the interaction of communities constructed from primary PPI network. The subcommunities constructed from all communities in the first level of organization constituted second level of organization. In the same way, successive levels were constructed until the level of *motifs,* thereby each smaller community had a minimum of one triangular *motif* defined by subgraph *G*(3,3). Since the triangular motif was overrepresented in PPI network and served as controlling unit in a network (19), we used *motif G*(3,3) as qualifying criteria for a community/subcommunity as a constituting member at a certain level of organization. Further, each community or smaller community landed up to different level of organization.

### Topological analyses of the networks

Cytoscape plugins, NetworkAnalyzer (20) and CytoNCA (21) were used to analyse the topological properties of the network for centralities, degree distribution, clustering coefficients and neighbourhood connectivity. The highest degree nodes were identified as *hubs* of the PCa network. Top 103 *hub* proteins having degree ***k*** *≥* 65 were considered for tracing the key regulators of the network. Other topological parameters, *viz.,* Rich club coefficients (***Φ***), Participation coefficients (***P_i_***) and Within-module degree (***Z_i_*** score) were calculated using Igraph package “brainGraph” (https://github.com/cwatson/brainGraph) in R. Another parameter subgraph centrality was calculated using Igraph functions.

#### Degree *(k)*

In the analysis of network, degree ***k*** indicates the total number of links established by a node in a network and is used to measure the local significance of a node in regulating the network. In a graph represented by *G* = (*N*, *E*), where *N* denotes nodes and *E* the edges, the degree of *i^th^* node (*k_i_*) is expressed as *k_i_* = ∑*_ij_^N^ A_ij_*, where *A_ij_* denotes the adjacency matrix elements of the graph.

#### Probability of degree distribution *P*(*k*)

It is the probability of a random node to have a degree ***k*** out of the total number of nodes in the network and is represented as fraction of nodes having degree (***k***), as shown in **Equation (1)**, where *N_k_* is the total number of nodes with degree *k* and *N*, total nodes in the network.

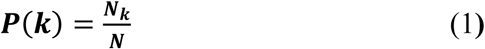

P(k) of random and small-world networks follow **Poison distribution** in degree distribution against degree, but most real-world networks, scale-free and hierarchical networks follow power law distribution *P(k) ∼ k^−γ^, where, 4 ≥ γ ≥ 2.* In hierarchical networks, *γ ∼* 2.26 (mean-field value) indicating a modular organization at different topological levels (22,23,24). Therefore, probability of degree distribution pattern defines the characteristic topology of a network.

#### Clustering coefficients *C(k)*

The strength of internal connectivity among the nodes neighbourhoods which quantifies the inherent clustering tendency of the nodes in the network is characterised by the Clustering coefficient ***C(k)***, which is the ratio between the number of triangular motifs formed by a node with its nearest neighbours and the maximum possible number of triangular motifs in the network. For any node *i* having degree *k_i_* in an undirected graph, ***C(k)*** can be expressed as **Equation (2)**, where *m_i_* is the total number of edges among its nearest-neighbours. In scale-free networks ***C(k)*** ∼ *constant*, but it exhibit power law in hierarchical network against degree, ***C(k)*** *∼ k*^−α^, with α ∼ 1 (22,23,24).

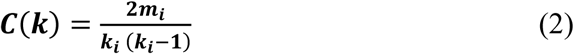

#### Neighbourhood connectivity *C_N_(k)*

The node neighbourhood connectivity (average connectivity established by the nearest-neighbours of a node with degree ***k***, represented by ***C_N_(k)*** can be expressed as shown in **Equation (3)**, where, *P(q|k)* is conditional probability of the links of a node with *k* connections to another node having *q* connections.

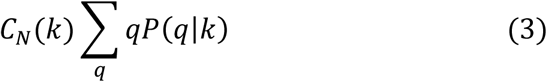

In hierarchical network topology, ***C_N_(k)*** exhibit power law against degree *k*, that is, ***C_N_(k)*** *∼ k^β^, where, β* ∼0.5 (25). Further, the *positivity* or *negativity of the exponent β* can be defined as, respectively, the assortivity or disassortivity nature of a network topology (26).

#### Centrality measures

A node’s global functional significance in regulating a network through information processing is estimated by the basic Centrality measures—Closeness centrality ***C_C_***, Betweenness centrality ***C_B_*** and Eigenvector centrality ***C_E_***(27). Another centrality measure, Subgraph centrality ***C_S_*** is also used to describe the participation of nodes in other subgraphs in the network (28). These centrality measures collectively determine the cost effectiveness and efficiency of information processing in a network.

The closeness centrality ***C_C_*** represents the total geodesic distance from a given node to all its other connected nodes. It represents the speed of spreading of information in a network from a node to other connected nodes (29). ***C_C_*** of a node *i* in a network is calculated by the division of total number of nodes in the network, *n* by sum of geodesic path lengths between nodes *i* and *j* which is represented *by d_ij_* in **Equation (4)**.

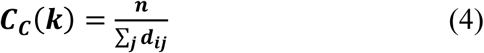

Betweenness Centrality ***C_B_*** is the measure of a node which is the share of all shortest-path traffic from all possible routes through nodes *i* to *j*. Thus, it characterizes a node’s ability to benefit extraction from the information flow in the network (30) and its controlling ability of signal processing over other nodes in the network (31,32). If *d_ij_*(*ν*) denotes the number of geodesic paths from node *i* to node *j* passing through node *ν*, then *C_B_*(*ν*) of node *ν* can be obtained by **Equation (5)**.

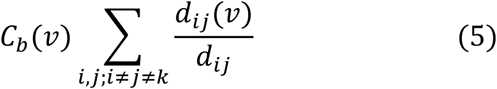

If *M* denotes the number of node pairs, excluding *v*, then normalized betweenness centrality is given by the **Equation (6)**.

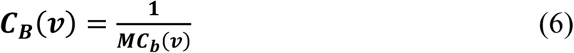

Eigenvector centrality ***C_E_*** is proportional to the sum of the centrality of all neighbours of a node and it reflects the intensity of these most prominent nodes influencing the signal processing in the network (33). If nearest neighbours of node *i* in the network is denoted by *nn(i)* with eigenvalue *λ* and eigenvector *v_i_* of eigen-value equations, *Aν_i_* = *λν_i_*(*ν*) where, *A* is the network adjacency matrix, ***C_E_*** can be shown by the **Equation (7)**,

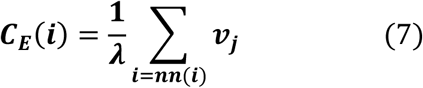

***C_E_*** score corresponds to maximum positive eigenvalue, *λ_max_*, of the principal eigenvector of *A* (34). Since a node’s ***C_E_*** function depends on the centralities of its neighbours, it varies across different networks association of high ***C_E_*** nodes; within closely connected locality of such nodes reduces the chances of isolation of nodes (33). Thus, ***C_E_*** becomes a powerful indicator of information transmission power of a node in the network.

The subgraph centrality ***C_S_*** of a node calculates the number of subgraphs the node participates in a network. It can be calculated using eigenvalues and eigenvectors of adjacency matrix of the graph, as shown in **Equation (8)**, where *λ_j_* is the *j^th^* eigenvalue and *v_j_(i),* the *i^th^* element of the associated eigenvector. The weightages are higher for smaller graphs. Higher subgraph centrality of a node corresponds to better efficiency of information transmission and increase in essentiality of the node in the network (28,35).

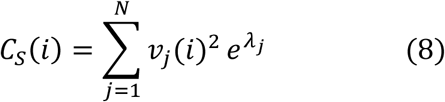

### Within-module degree and Participation coefficients of the *hubs*

In complex networks the characterization of hubs as high degree nodes with higher centrality values is incomplete without exploring the role of nodes at the modular levels (36). The role of nodes at the modular level is determined through the participation of nodes in establishing links between the nodes within the module as well as outside the module and calculating the modular degree of the nodes. Within-module degree or Z-score, ***Z_i_***, signifies the connections of a node i in the modules and categorizes a node as modular hub-node with high (***Z_i_*** ≥ 2.5) signifying more intra-module connectivity of the node than inter-module, whereas, lower *Z* values, ***Z_i_*** < 2.5, categorizes as non-*hub* nodes with less intra-module connectivity (36). The *Z-*score can be calculated as shown in **Equation (9)**, where *k_i_* represents the number of links of node *i* to other nodes in its modules *s_i_* and *k̄_s_i__* the average of degree ***(k)*** over all nodes in *s_i_*; *σk̄_s_i__* is the standard deviation of *k* in *s_i_*.

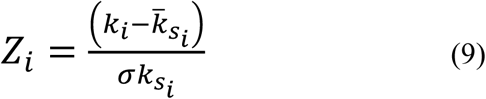

The participation coefficient, ***P_i_***determines the participation of the node *i* in linking the nodes inside and outside its module (36). ***P_i_***values lie in the range of *0-1* with higher values corresponding to the participation of nodes in establishing links outside the modules with homogeneous distribution of its links among all modules, and if *k_is_* is taken to represent the number of links of node *i* to nodes in modules *s* and *k_i_*, the total degree of node *i*, ***P_i_*** can be calculated as in **Equation (10)**, where, N_M_ is the number of modules in the network.

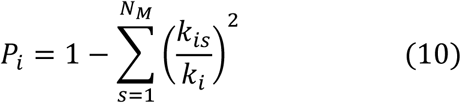

### Rich-club analysis

Identification of hubs in a network generally is done through general centrality measures, especially higher degree nodes are commonly considered as hubs and existence of high degree nodes in a network correlate with the local regulatory roles of these high degree hubs in the network (37). This phenomenon of formation of rich club connection between high degree hubs exhibit the robustness of the network and the resilience when the hubs are targeted (38). The existence of rich club phenomenon among hubs is investigated by calculating the Rich-club coefficients ***Φ(k)*** across the degree range (38). ***Φ(k)*** is equivalent to the clustering coefficient among a subgroup of nodes with degrees ≥ ***k***. In order to remove the random interconnection probability factor, normalization of the rich club coefficients can be done by the **Equation (11)**, where ***Φ_rand_(k)*** is the rich-club coefficient of random networks with similar size and degree sequence and ***Φ_norm_(k)*>1** indicating a rich-club formation. This rich club phenomenon is associated with the *assortivity* nature of the networks and is important to understand the roles played by these hubs roles in the network integration and efficient transmission of signals (39).

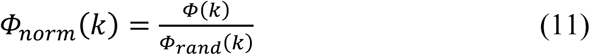

### Tracking the key regulators in the networks

The most influential genes in the PCa network was identified first through calculating the centrality measures. Since, higher degree nodes have higher centrality values, top 103 highest degree nodes (Degree ***k*** ≥ 65) were considered among the *hub* nodes of the network for tracing the key regulators which may play important role in regulating the network. Then tracing of nodes from the primary network up to *motif* level *G(3,3)* was done on the basis of representation of the respective nodes (proteins) across the sub modules obtained from Louvain method of community detection/ clustering. Finally, the *hub*-nodes (proteins) which were represented at the modules at every hierarchical level were considered as key regulators of the PCa network.

### Functional association analysis of modules

The modules at all levels of hierarchy were analysed for their functional annotations with DAVID functional annotation tool (40,41). The functions and pathways with corrected *p*<0.05 were considered statistically significant.

## Results

### PPI network in PCa follows hierarchical scale-free topology composed of modules at five levels of hierarchy

From the interactome network of 3,871 PCa genes, the physical interacting PPI network of 2,960 proteins with 2,960 nodes and 20,372 edges was constructed as the primary network (**Figure 1**). Analysis of this primary PCa network showed that the network followed power law distributions for probability of node degree distribution, *P(k),* clustering coefficient *C(k)* and neighbourhood connectivity distribution *C_N_(k)* against degree *(k)* with *negative exponents* (22) **Equation (12)** (**Figure 3**). This power law feature indicates that the network exhibited hierarchical-scale free behaviour with systems level organization of modules/communities. Further, community finding using Louvain modularity optimization method (18) led to the detection of communities and sub-communities at various levels of organization (**Figure 2A**). Thus, a total of 436 communities and smaller communities were detected, out of which 38 reached up to level **V**, the level of *motif G(3,3)*.

**Figure 1:**
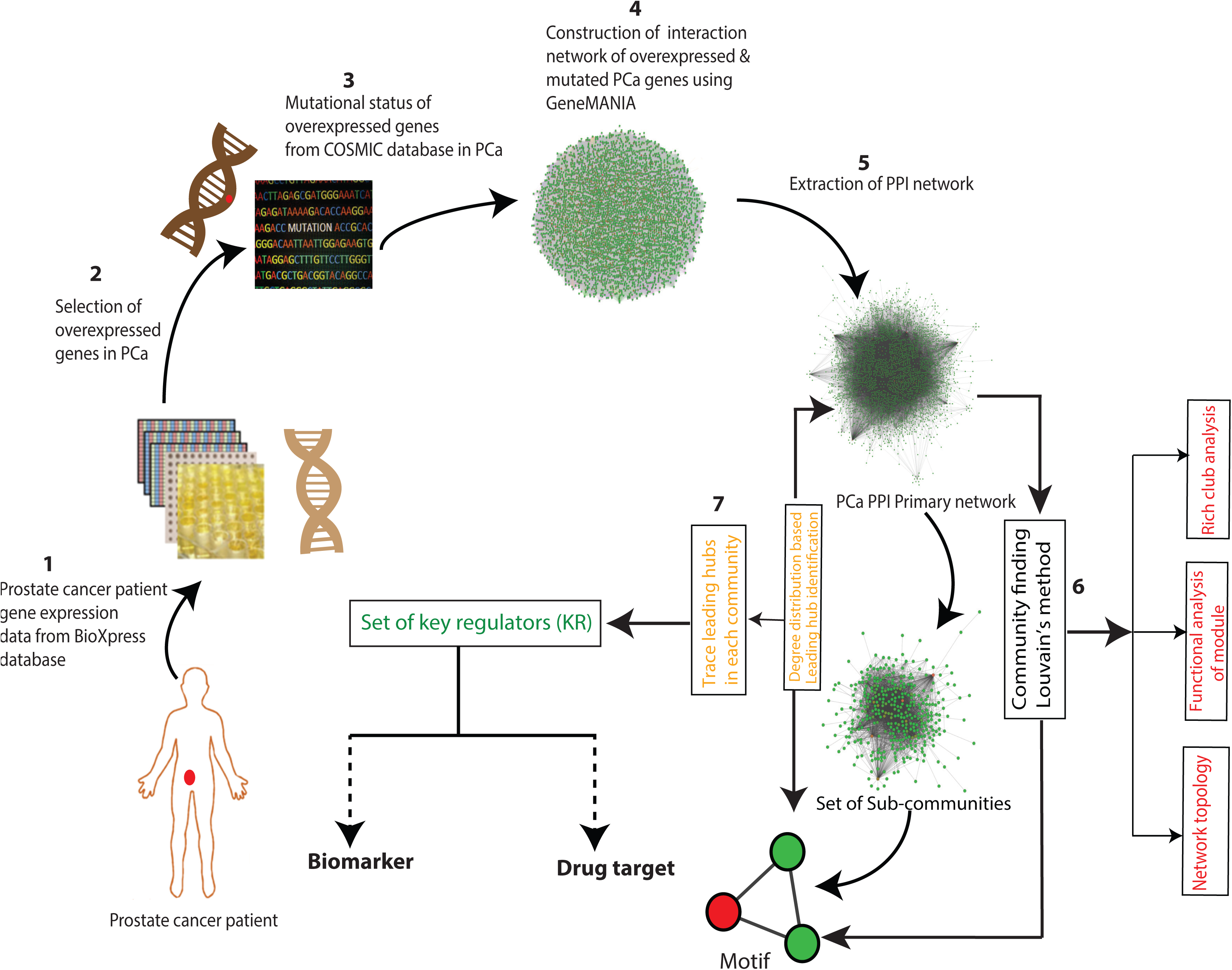
Flowchart of the methodology.

**Figure 2:**
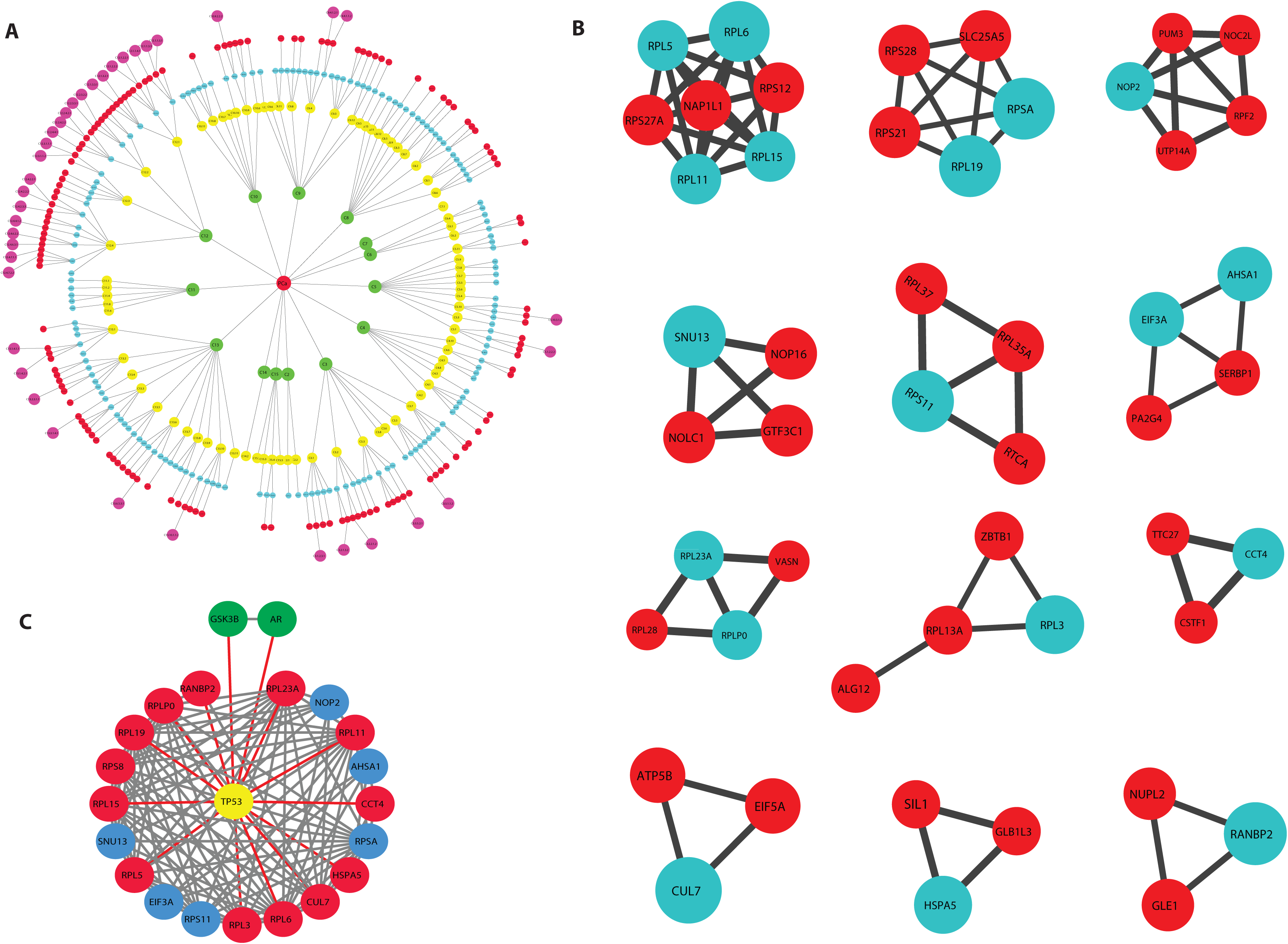
**A.** Communities/modules of PCa PPI network. **B.** Interacting partners of the 19 key regulators at motif level. **C.** Interaction of key regulators with p53, AR and GSKβ.

**Figure 3:**
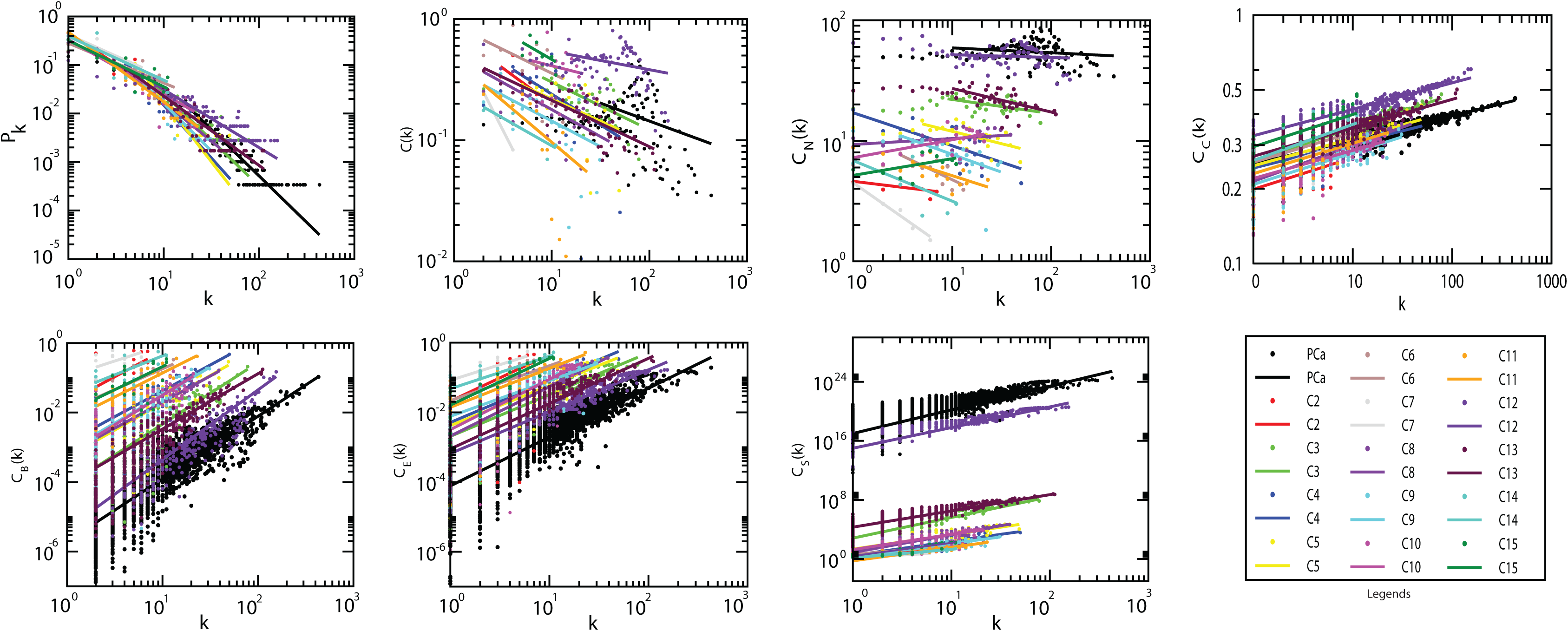
Topological properties of PCa and the modules/communities at the first hierarchical level. Degree distribution probability (*P*(*k*)), clustering coefficient (*C*(*k*)), neighbourhood connectivity (*C_N_*(*K*)) as function of degree (*k*) and centrality measurement closeness (*C_C_*(*k*)), betweenness centrality (*C_B_*(*k*)), eigenvector centrality ((*C_E_*(*k*))), subgraph centrality (*C_S_*) as a function of degree.

Communities at the first hierarchical level also showed power law distribution for ***P(k), C(k)*** and ***C_N_(k)*** against degree distribution with negative exponents indicating further systems level organization of modules (**Equation 12**) except in case of communities C8, C10 and C15 where the ***C_N_(k)*** exhibit power law against degree k with positive exponents (*β ∼ 0.05,0.13,0.14* respectively) (**Figure 3**). This indicates assortivity nature in the modules indicating the possibility of rich-club formation in these modules, where, hubs play significant role in maintaining network properties and stability (25).

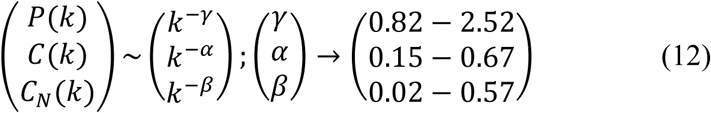

### Nineteen (19) novel regulators served as backbone of the network

Centrality measures are used to assess the importance of the nodes in information processing in a network. Betweenness centrality ***C_B_***, Closeness centrality ***C_C_***, Eigenvector centrality ***C_E_*** and Subgraph centrality ***C_S_***are various topological properties which can determine the efficiency of signal transmission in a network (28,42). In PCa network and modules at the first hierarchical level, these parameters also exhibited power law as a function of degree ***(k)*** with *positive exponents* where the centralities tend to increase with higher degree nodes (**Equation 13**) (**Figure 3**). This behaviour revealed the increase in efficiency of signal processing with higher degree nodes in the network showing the importance of these nodes in controlling the flow of information, thereby regulating and stabilizing the network. Hence, *hub* proteins had a significant influence in regulating the network and might be playing an important role in PCa. In order to identify the most influential key regulator proteins in the network, top 103 *hub*-proteins having degree *(**k**) ≥ 65* were considered for identification of the key regulators through their representation at every topological level (**Supplementary Table 1**). After tracing *hubs* at every topological level, 19 (RPL11, RPL15, RPL19, RPL23A, RPL3, RPL5, RPL6, RPLP0, RPS11, RPS8, RPSA, HSPA5, NOP2, RANBP2, SNU13, CUL7, CCT4, ASHA1 and EIF3A) (**Table 1 & 2**) were found to be the backbone of the network. These key regulators along with their partners forming the *motifs* (**Figure 2B**), might be playing the most important roles in regulating and maintaining the stability (network integrity, optimization of signal processing, dynamics etc) of the network.

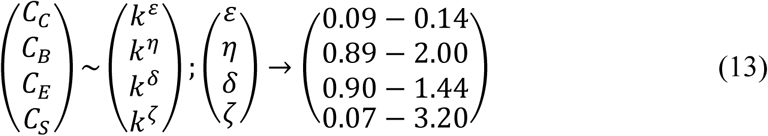

### Modules of the network were associated with specific functions

Community detection of the network using Louvain modularity optimization method leads to clustering of the primary PCa network up to the level of *motifs* (**Figure 2A**). This clustering showed that Modularity (***Q***) of the networks exhibited an increasing pattern with topological levels with highest average Modularity (***Q*** = 0.5527) seen at the first hierarchical level, and lowest (***Q*** = 0.0013) at the level V, the *motif* level (43,44).

In complex PPI network the modules have biological meanings relating to functions and gene ontology analyses have revealed enrichment of certain known functions and pathways in the modules (45). Our primary PCa-network was composed of 14 modules deduced from the community detection and their mean clustering coefficients ***C(k)*** ∼ 0.094 - 0.392 (**Table 3**). Among these, modules C12 and C13 which were the largest and had the highest mean clustering coefficients ***C(k)***=0.392 and 0.218, respectively, showing a functional homogeneity in the modules. These modules were analysed for their functional annotations with DAVID functional annotation tool (40,41) to reveal association with different functions (**Table 3**).

**Table 1.** Key regulators and their topological properties.

| Sl. No. | ID | Gene | Function | Degree ($k$) | Closeness centrality ($C_C$) | Betweenness centrality ($C_B$) | Eigenvector centrality ($C_E$) | Subgraph centrality ($C_S$) |
| --- | --- | --- | --- | --- | --- | --- | --- | --- |
| 1. | *AHSA1* | Activator of Hsp90 ATPase activity 1 | Positive regulation of ATPase | 75 | 0.379262 | 0.00331 | 0.054588 | 2.47E+23 |
| 2. | *CCT4* | Chaperonin containing TCP1 subunit 4 | Protein folding | 66 | 0.384435 | 0.003334 | 0.033523 | 9.29E+22 |
| 3. | *CUL7* | Cullin 7 | Ubiquitin-dependent protein catabolism | 270 | 0.435916 | 0.034463 | 0.151248 | 1.89E+24 |
| 4. | *EIF3A* | Eukaryotic translation initiation factor 3 subunit A | Translation pre-initiation complex formation | 85 | 0.37006 | 0.002209 | 0.079073 | 5.18E+23 |
| 5. | *HSPA5* | Heat shock protein family A (Hsp70) member 5 | Activation of signaling protein activity involved in unfolded protein response | 111 | 0.408758 | 0.012349 | 0.062591 | 3.24E+23 |
| 6. | *NOP2* | NOP2 nucleolar protein | rRNA base methylation, | 68 | 0.375317 | 0.001319 | 0.076741 | 4.88E+23 |
| 7. | *RANBP2* | RAN binding protein 2 | Protein sumoylation | 72 | 0.386242 | 0.004881 | 0.031476 | 8.19E+22 |
| 8. | *RPL11* | Ribosomal protein L11 | Ribosomal large subunit assembly | 83 | 0.383638 | 0.001239 | 0.114333 | 1.08E+24 |
| 9. | *RPL15* | Ribosomal protein L15 | Nuclear-transcribed mRNA catabolic process | 79 | 0.382201 | 0.001289 | 0.109354 | 9.91E+23 |
| 10 | *RPL19* | Ribosomal protein L19 | Nuclear-transcribed mRNA catabolic process | 78 | 0.381856 | 0.000469 | 0.113259 | 1.06E+24 |
| 11 | *RPL23A* | Ribosomal protein L23a | Ribosomal large subunit assembly | 84 | 0.379602 | 0.001182 | 0.109817 | 9.99E+23 |
| 12 | *RPL3* | Ribosomal protein L3 | Ribosomal large subunit assembly | 67 | 0.385588 | 0.000788 | 0.100707 | 8.41E+23 |
| 13 | *RPL5* | Ribosomal protein L5 | Ribosomal large subunit assembly | 92 | 0.383141 | 0.001438 | 0.114815 | 1.09E+24 |
| 14 | *RPL6* | Ribosomal protein L6 | Ribosomal large subunit assembly | 113 | 0.39809 | 0.003601 | 0.129049 | 1.38E+24 |
| 15 | *RPLP0* | Ribosomal protein lateral stalk subunit P0 | Nuclear-transcribed mRNA catabolic process | 88 | 0.394955 | 0.002486 | 0.110764 | 1.02E+24 |
| 16 | *RPS11* | Ribosomal protein S11 | Nuclear-transcribed mRNA catabolic process | 74 | 0.376943 | 0.000751 | 0.102129 | 8.64E+23 |
| 17 | *RPS8* | Ribosomal protein S8 | Nuclear-transcribed mRNA catabolic process | 120 | 0.394218 | 0.004481 | 0.126136 | 1.32E+24 |
| 18 | *RPSA* | Ribosomal protein SA | Ribosomal small subunit assembly | 79 | 0.378002 | 0.002864 | 0.094957 | 7.47E+23 |
| 19 | *SNU13* | SNU13 homolog, small nuclear ribonucleoprotein (U4/U6, U5) | mRNA splicing, via spliceosome | 87 | 0.370245 | 0.003088 | 0.072649 | 4.37E+23 |

**Table 2.**
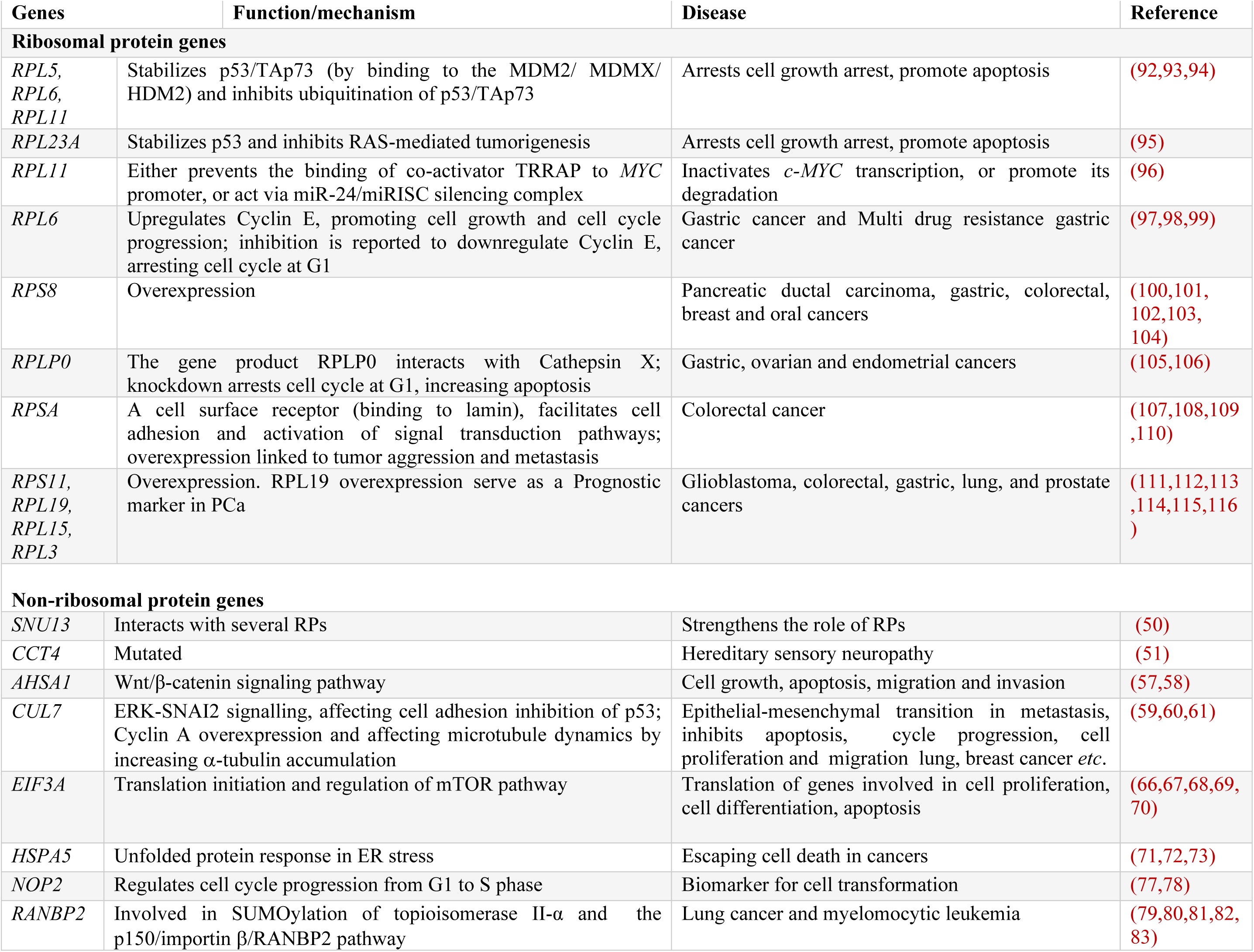
The key regulators identified in this study and their key functions in disease condition.

**Table 3.** Average Clustering coefficients of the PCa modules at first hierarchical level.

| **Sl. No.** | **Modules** | **Most enriched function** | **Avg. Clustering coefficient** | **Corrected *p*-value** |
| --- | --- | --- | --- | --- |
| 1. | C2 | RNA mediated gene silencing | 0.114 | 2.07E-07 |
| 2. | C3 | Unfolded protein folding | 0.209 | 4.12E-16 |
| 3. | C4 | ATP/nucleotide binding | 0.171 | 2.64E-14 |
| 4. | C5 | MRNA transport | 0.165 | 2.25E-15 |
| 5. | C6 | Transcription initiation | 0.374 | 2.68E-12 |
| 6. | C7 | Endoplasmic reticulum membrane proteins | 0.094 | 0.019 |
| 7. | C8 | Endocytosis | 0.14 | 2.67E-09 |
| 8. | C9 | Mitochondrial proteins | 0.124 | 6.75E-33 |
| 9. | C10 | Proteosome | 0.182 | 1.02E-16 |
| 10. | C11 | Ubiquitin protein ligase activity | 0.107 | 1.13E-10 |
| 11. | C12 | Ribonucleoprotein | 0.392 | 7.50E-104 |
| 12. | C13 | Transcription regulation | 0.218 | 6.11E-74 |
| 13. | C14 | Transmembrane helix | 0.096 | 4.07E-04 |
| 14. | C15 | DNA repair | 0.31 | 6.61E-08 |

### Hubs in the PCa network coordinate the modules acting as modular hubs

In complex hierarchical networks, the modularity of sub-communities and the roles played by the nodes in the modules is defined with the nodes Within-module Z score*, **Z_i_*** along with their Participation coefficients ***P_i_*** (36). ***Z_i_*** gives the degree of the nodes within their modules, and ***P_i_*** describes the influence of a node inside the module, as well outside it, in terms of signal processing as well as maintaining network stabilization. Hence, ***Z_i_***and ***P_i_*** were calculated for each node in the modules using **Equations (9) & (10)**, respectively. Accordingly, within-module Z score, the nodes are classified as follows:

1. **Modular non-hub nodes *Z_i_ < 2.5:*** *(R1) Ultraperipheral nodes*: The nodes linking all other nodes within their modules, ***P_i_*** ≤ 0.05 *(R2) Peripheral nodes:* nodes linking most other nodes in their module, 0.05 < ***P_i_*** ≤ 0.62; *(R3) non-hub connector nodes*: nodes linking many nodes in other modules, 0.62 < ***P_i_*** ≤ 0.80; and *(R4) Non-hub kinless nodes*: nodes linking all other modules, ***P_i_***> 0.80.
2. **Modular hubs *Z_i_ > 2.5:*** *(R5) Provincial hubs*; hub nodes linking vast majority nodes within their modules, ***P_i_*** ≤ 0.30; *(R6) Connector hubs*; hubs linking most the other modules, 0.30 < ***P_i_***≤0.75; and *(R7) Kinless hubs*; hubs linking among all modules, ***P_i_*** > 0.75.

In the PCa network in this study, many *hub*-proteins were acting as *modular hubs,* helping in establishing connection between the modules at different hierarchical levels. For example, CUL7 and RANBP2 were among important key regulator protein *hubs* in PCa which also acted as *modular kinless* and *connector hubs* of module C3 and C5 at the first hierarchical level (**Figure 4B**). P53, E2F1 and c-MYC acted as *kinless global hubs* of module C9 connecting with all the modules and other proteins in the network. NOP56, FBL, RNF2 and NPM1 also acted as *connector modular hubs* of module C12, connecting other modules at the same level (**Figure 4A** **& 4C**).

**Figure 4:**
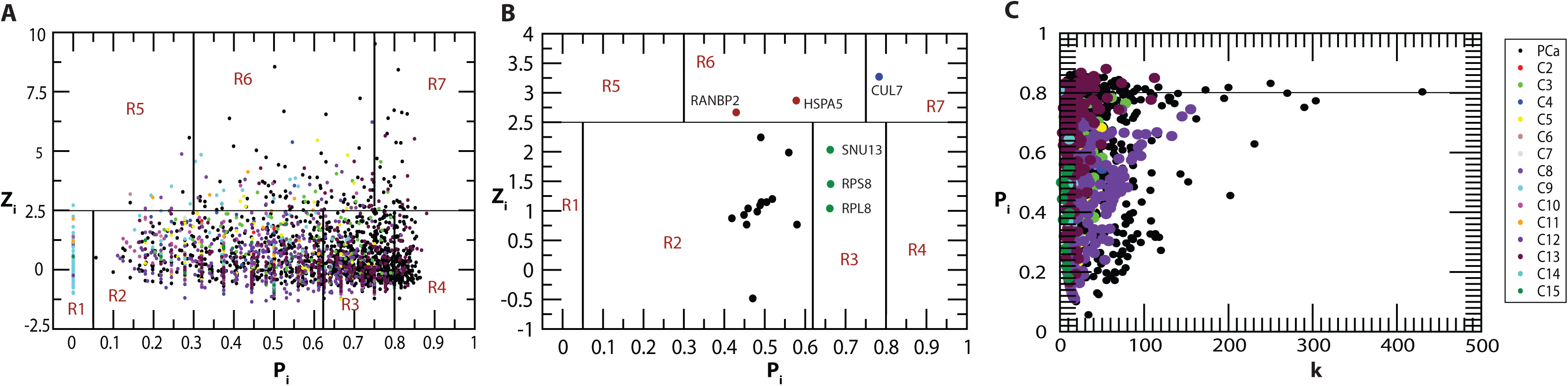
Identification of modular hubs **A.** In the primary PCa network and the modules at first Hierarchical level with within module *Z* score *Z_i_*, and their participation coefficients P_i_. **B.** Identification of modular hubs among 19 key regulators. **C.** Participation coefficient vs degree in PCa primary network and the modules at first hierarchical level.

### PCa network exhibited non-monotonicity in rich-club formation across the hierarchy

Identification of rich club nodes is another common feature to study the influence of hubs in the network forming a strong connection among them which is done by calculating normalized rich club coefficient ***Φ_norm_*** across the degree range ***k*** (**Equation 11**). Normalized rich-club coefficient ***Φ_norm_***> 1 indicates the existence of rich club among the nodes which play key role in network integration, increasing its stability and improving the efficiency of transmission of information among hub proteins. Since, PCa network is hierarchical and shows *disassortativity* in nature with node neighbourhood connectivity ***C_N_(k)*** following power law distribution against degree *(**k**)* with *negative value of exponent β* (**Equation 13**), rich club formation among the hub proteins is quite unlikely (38,46). Although rich club formation is not exhibited among high degree *hub* proteins, the moderate intermediate degree protein with degree ***(k)*** ∼ 19 - 104 showed higher rich club coefficients than the *hubs* in PCa network (**Figure 5**). In the PCa network across the hierarchy, different patterns of rich club coefficients were exhibited among the modules (**Figure 5**), showing the phenomenon of non-monotonic behaviour at different hierarchical levels. With respect to modules C12 and C13 at first hierarchical level, they exhibit rich club formation between the high degree nodes but the pattern changes moving at the lower levels. However, in the modules *C*8, *C*10 and *C*15, the topological properties of these modules exhibit *assortativity* nature due to (i) the node neighbourhood connectivity ***C_N_***(***k***) in these modules follow power law with positive *β* exponents, (ii) ***Φ*** increases monotonically with degree k, and (iii) ***Φ_norm_*** approximately increases with degree ***k*** with values of ***Φ_norm_*** > 1 (**Figure 6**), indicating the possibility of rich club formation among the high degree nodes (**Figure 6A**). Considering the nodes with degrees whose ***Φ_norm_*** is larger than one, the approximate range of degrees of nodes forming rich-club in these three modules are 61 *≥ k ≥* 14 (*C*8), 52 *≥ k ≥* 6 (*C*10), 37*≥ k ≥* 6 (*C*15), and clearly show rich-club formations in the respective network modules (red coloured nodes in the respective modules in **Figure 6**.

**Figure 5:**
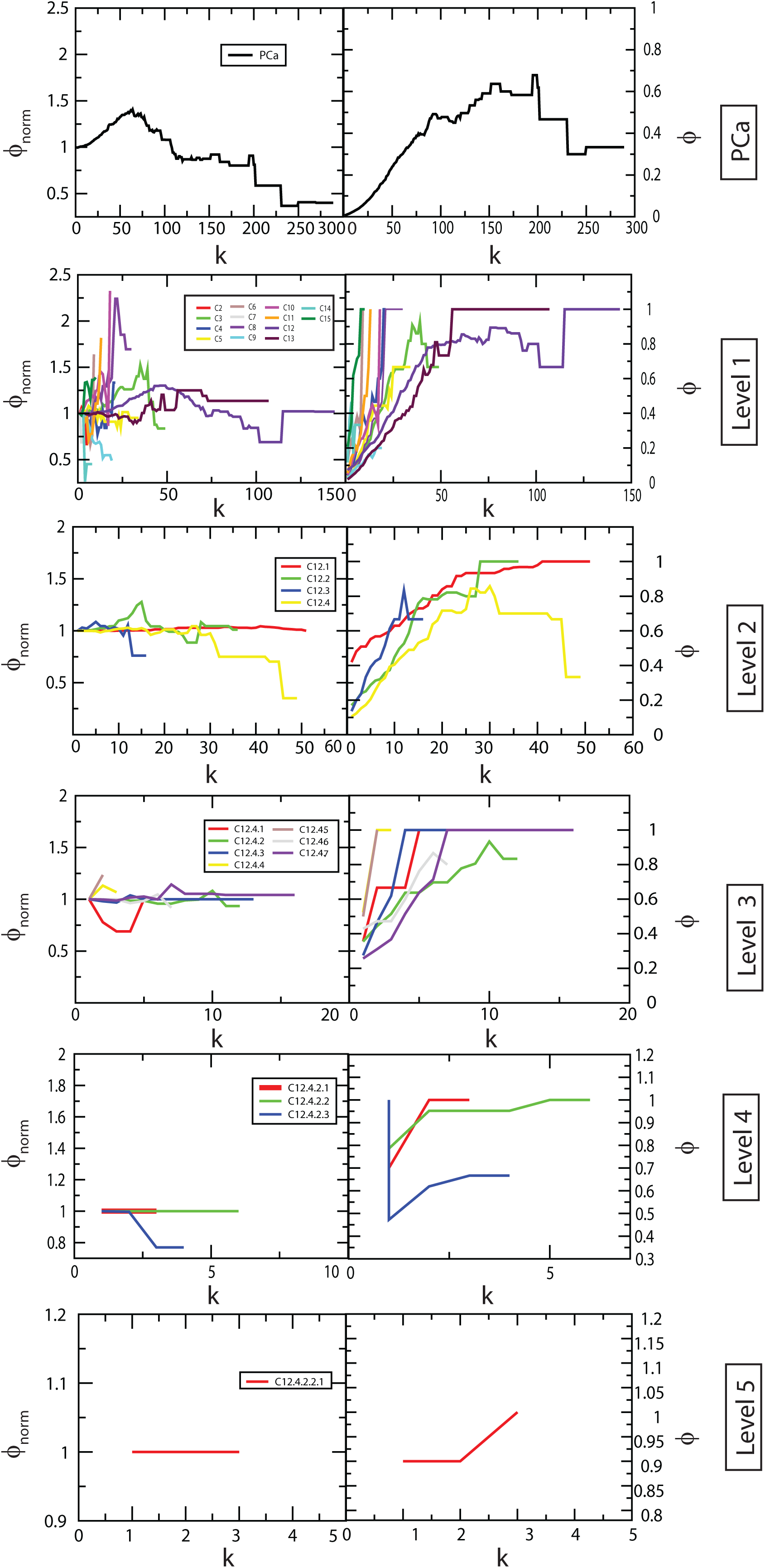
Rich club analysis of PCa PPI network and the communities upto the last motif level.

**Figure 6:**
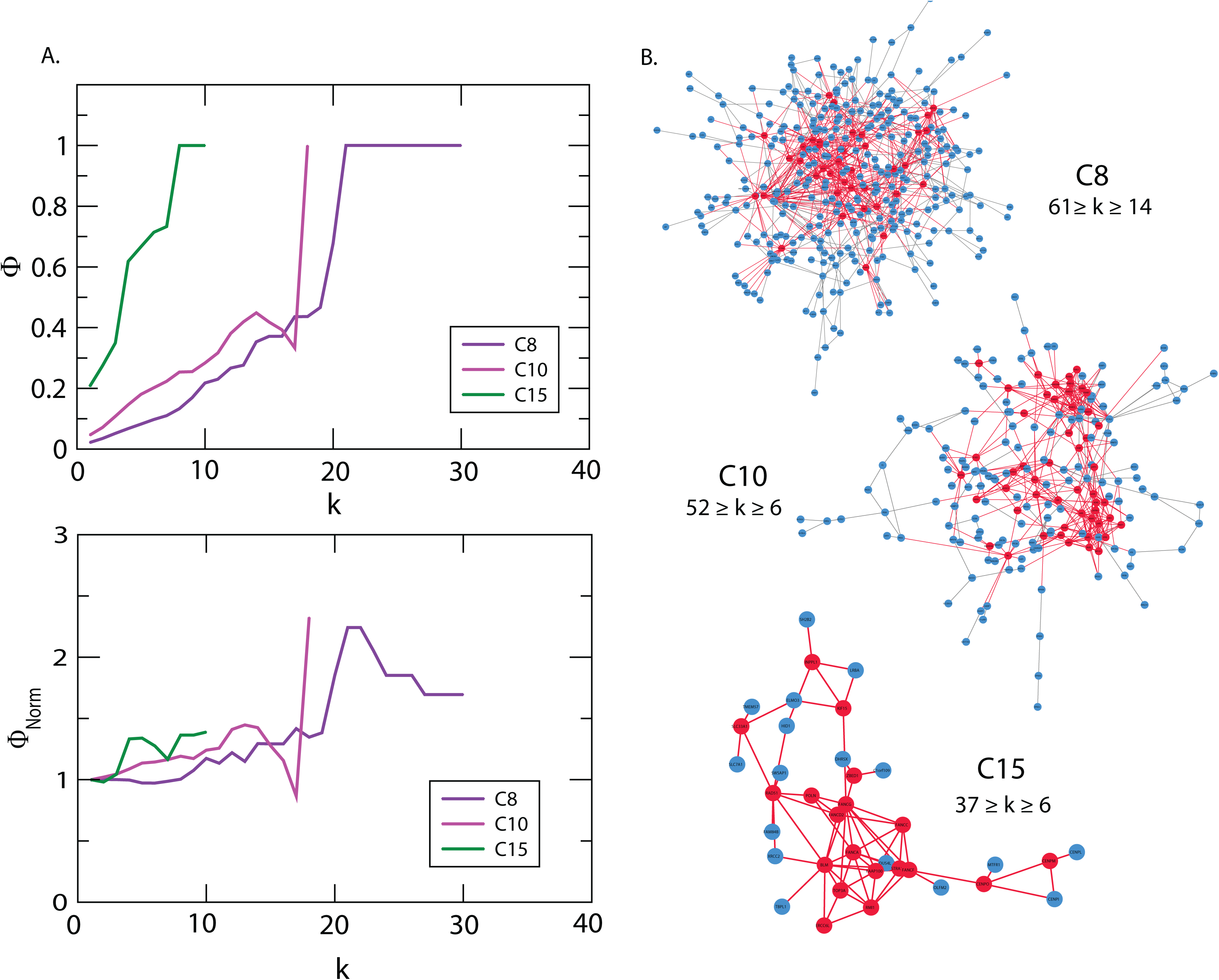
**A.** Rich formation in C8, C10 and C15 in first hierarchal level of PCa. **B.** Degree maximum and minimum degrees of rich club forming hubs.

## Discussion

The real-world complex networks generally have hierarchically organized community structure, which is evident from fractal studies and scaling behaviour of these networks (22). Even though there is no specific definition of communities or modules in a network, each community/module is established by densely interconnected nodes forming clusters around the hub nodes which generally have their own local properties and organization (43). The hubs have highest interactions in the network due to their high-degree, constitute both intra- and inter-communities’ interactions in the network in a hierarchical manner, and thus play a central role in information processing in the network (37). The primary PPI PCa network constructed in this study for tracking the hubs up to the level of *motifs* led to the identification of 19 key regulators (*hubs*) from 3,871 genes found to be significantly overexpressed in human prostate adenocarcinomas.

There have been limited community finding methods in complex networks, among which the Newman and Girvan leading eigenvector algorithm (47), is commonly used. However, in comparatively large complex networks, Louvain method, which is based on modularity, ***Q*** maximization/optimization (18), is the most suitable, sensitive and comparatively faster. In our study, considering the size of the network and its sensitivity, we used Louvain method for community detection and while giving equal importance to the *hubs*, *motifs* and *modules* of the network, we identified the novel key regulators. 11 key regulators (*RPL*11, *RPL*15, *RPL*19, *RPL*23A, *RPL*3, *RPL*5, *RPL*6, *RPL*P0, *RPS*11, *RPS*8 and *RPSA*) belong to the family of ribosomal proteins (RPs) which are involved in ribosomal biosynthesis and other eight predicted regulators (*HSPA*5, *NOP*2, *RANBP*2, *SNU*13, *CUL*7, *CCT*4, *ASHA*1, *EIF3A*) have other important functions and also reported to be associated with various other cancers. Moreover, at the level of *motifs* these key regulators interact with other proteins which may also be playing important roles in PCa and establishing themselves to be the candidate disease-genes along with key PCa regulators (**Figure 2B**).

The emergence of 11 RPs as key regulators in PCa is an important finding in this study. It could be due to the crucial role of RPs in cell growth and proliferation propagated through protein synthesis. In cancers, ribosomal biosynthesis increases to meet the requirement of rapidly growing/proliferating cells (48). Some RPs take part in extra-ribosomal functions involved in tumorigenesis, immune cell signalling, and development and regulating diseases through translocation across the nuclear pore complex (49). RPs have been associated with tumorigenesis either as oncoproteins or tumour suppressors, with differential roles being reported in different cancers. During ribosomal or nucleolar stress such as hypoxia, lack of nutrient, starvation, deregulation of genes *etc*., RPs modulate the p53-mediated apoptosis. The association of RPs with cancers as discussed in **Table 2** suggests a potential unexplored function of these proteins in PCa, both as therapeutic target and as predictive biomarker. An understanding of the functions and the pathways of key RPs, for example their role in stabilizing p53 during ribosomal stress and role in cell growth/proliferation in PCa patients is of immense significance as it provides new insights into the control and prevention of PCa.

Besides, other non-ribosomal predicted key regulators identified in this study, *SNU*13, *CCT*4, *AHSA*1, *CUL*7, *EIF*3*A*, *HSPA*5, *NOP*2 and *RANBP*2, are also vital in cell physiology and are equally important for their involvement in cell growth and proliferation in one way or another. The NHP2-like protein 1 (SNU13) identified in this study as another key regulator, is a component of the spliceosome complex (50) which interacts with several RPs and strengthens the role of RPs in cancers. CCT4, Chaperonin containing TCP1 subunit 4, is a chaperone which when mutated is associated with hereditary sensory neuropathy (51).

AHSA1, the Activator of HSP90 ATPase Activity 1, is a positive regulator of the heat shock protein 90 (HSP90) (52). The activated HSP90 forms a complex with HSP70 and helps in either binding of the tumour suppressor p53 to DNA, or its degradation by ubiquitination (53). In cancers, activated HSP90 stabilizes the mutated p53 which decreases its DNA binding activity and degradation through binding with its inhibitor MDM2, thus promoting tumour progression (54,55). The activation and transportation of steroid hormones (androgen receptor, AR and oestrogen receptor, ER) to the nucleus is also mediated by HSP90 (56); thus, AHSA1 activation of HSP90 may influence the androgen metabolism in PCa. Moreover, AHSA1 is a regulator of the cell growth, apoptosis, migration and invasion through Wnt/β-catenin signaling pathway (57,58), which suggests its role as a candidate-disease gene in PCa.

CUL7, Culin7, is a component of an E3 ubiquitin-protein ligase complex and interacts with p53, CUL9 and FBXW8, and is reported to be an antiapoptotic oncogene (59). CUL7 has been associated with various cancer types, but its promotion of epithelial-mesenchymal transition in metastasis and its regulation of ERK-SNAI2 signalling affecting the expression of cell adhesion proteins, E-cadherins, fibronectin, N-cadherin and vimentin in cancer is well studied (60). CUL7 inhibits apoptosis in lung cancer through inhibition of p53 which regulates c-MYC cell cycle progression (59). CUL7 regulates cell cycle progression through Cyclin A overexpression, and also affects the cell migration, which is a hallmark of cancer, affecting microtubule dynamics in breast cancer (61). Therefore, the targeted knockdown and silencing of CUL7 has led to a decrease in cell proliferation, weaker α-tubulin accumulation in microtubules, promoting their stability and decreasing cell migration (in breast, liver and lung carcinoma cells) and has been suggested as a potential therapeutic target in various cancers (59,61,62,63).

The Eukaryotic translation initiation factor 3 subunit A (EIF3A) forms 43S Pre-initiation complex (43S PIC) with other initiation factors and 40S ribosome and initiates the protein synthesis process. This translates mainly genes involved in cell proliferation, cell differentiation, apoptosis *etc*. and exerts transcriptional activation/repression through forming different forms of stem loop binding with the mRNAs (64, 65). Dysregulation of translation initiation and the role of EIF3 has been studied in cancers (66,67,68). Moreover, involvement of EIF3 complex in regulation of mTOR pathway, which is associated with many cancers (69,70), makes it an interesting protein to study for its regulatory role in PCa.

The Heat shock protein family A (HSP70) member 5 (HSPA5) or glucose-regulated protein 78 kDa (GRP78), is a chaperone localized in endoplasmic reticulum (ER) and involved in folding and assembly of proteins and plays an active role in unfolded protein response in ER stress, promoting cell survival which is a common process of escaping cell death in cancers (71,72,73). Due to this activity, HSPA5 is an emerging therapeutic drug target for cancer.

NOP2 (p120) is a putative RNA methyl transferase protein and its expression is detectable in proliferating normal and tumour cells, but undetectable in non-proliferating normal cells (74,75,76). Its role in regulating cell cycle progression from G1 to S phase and transformation of normal fibroblast cells (77,78) makes NOP2 an interesting protein which can be used as biomarker for cell transformation. The Ran binding protein 2 (RANBP2) is another key regulator identified in this study which is involved in the SUMOylation of Topioisomerase II-α before the onset of anaphase, helping in separation of chromatids from the centromere and its under-expression, mutation or deficiency has been observed in various cancers specially lung cancer and myelomocytic leukemia acting as tumor suppressor genes (79,80,81,82,83). Since SUMOylation plays an important role in tumour progression (84), the p150/importin β/RANBP2 pathway may also play a significant role in PCa progression.

In PCa, p53 and AR are the most mutated genes reported according to COSMIC (15). Association of mutation in the androgen receptor gene (AR) which causes the mutated receptor to be always in activated state and continue to maintain androgen receptor mediated downstream signalling even in lower level of circulating androgens leading to discovery of androgen independency in prostate cancer (85). A recent report suggests several mutations in the AR gene in different metastatic castration-resistance (CRPC) patients in prostate cancer suggesting AR mutants as a good biomarker candidate (86). *β*-catenin and GSK-3β are other co-regulators of Androgen receptor and phosphorylation of AR by GSK-3β which inhibit AR driven transcription, but in prostate cancer, the increase in the activity of Akt suppression of GSK-3β due to phosphorylation helps in PCa progression (87,88). Out of the 19 key regulators identified in this study, 13 (*CUL*7, *HSPA*5, *CCT*4, *RPL*19, *RPL*11, *RPL*3, *RPL*6, *RPLP*0, *RPL*5, *RANBP*2, *RPS*8, *RPL*23*A* and *RPL*15) interact directly with p53 and other key regulators through them (**Figure 2C**). In the PCa network, AR interacted with these key regulators through p53 and GSK-3β, which is its upstream regulator (87,88). The observations suggest an important regulatory role of the reported key regulators in regulating the functions mediated through p53 and AR in PCa. The findings reiterate the putative roles of these *hubs* in PCa manifestation and progression. This study may prove fundamental in characterizing the potential therapeutic targets and biomarkers for sensitive intervention and diagnosis of PCa.

It is to be noted that in this study the PCa PPI network follows Hierarchical scale free topology. Along with the conventional centrality measures, *C_B_,C_C_,C_E_* and *C_S_,* probability degree distribution *P(k),* clustering coefficient *C(k)* and node neighbourhood connectivity distribution *C_N_(k)* are used to characterize a network whether one is scale-free, random, small-network or hierarchical network (22,24,42). PCa PPI network follows power law distributions for probability of node degree distribution, *P(k)*, clustering coefficient, *C(k)*, and neighbourhood connectivity distribution against degree *k* with *negative exponents* (22) (**Equation 12**) (**Figure 3**), indicating the network falls in hierarchical-scale free behaviour which can exhibit systems level organization of modules/communities.

Since, node neighbourhood connectivity distribution *C_N_(k)* as a function of degree *k* obeys power law with negative exponent *β*, it shows its *disassortative* nature indicating that there is no signature of rich club formation among high degree nodes in the network (38). Degree centrality is the most commonly used centrality measure used to define the *hubs* which are the high degree nodes in the network. This *disassortivity* may be due to the sparse distribution of the *hubs* among the modules playing key roles in coordinating specific function within each module as well as establishing the connections among the modules (38). Furthermore, we used Louvain modularity optimization method (18) to detect, find communities and subcommunities and their organization at various levels of organization (**Figure 2A**). The communities/subcommunities at various hierarchically organized levels also exhibited hierarchical scale-free topology, as was the case in the primary PCa network (**Figure 3**). This hierarchical organization shows the systematic coordinating role of the emerged modules/communities and *hubs* in regulating and maintaining the properties of the network (10). In such type of networks, the centrality-lethality rule (37) is not obeyed which indicates that disturbing the hub/hubs in the network will not cause the whole network collapse.

Another important feature we found in PCa network is the observation of the nonmonotonic behaviour in the rich club formation in the PCa PPI network and across its hierarchy (**Figure 5**). The intermediate nodes in PCa network shows higher rich club coefficients than the highest degree hubs, indicating an important role of these intermediate nodes in regulating the network organization and maintaining stability through formation of key links between the low degree nodes and high degree hubs. Formation of rich club among the high degree nodes in the communities C8, C10 and C15 (**Figure 6A**) indicating an increase in sensitivity of these *hubs* on being targeted hence take significant roles in regulating in their respective modular functions, i.e., endocytosis, proteosome and DNA repair mechanisms (**Table 3**). These high degree hubs in these modules fall among the intermediate degree nodes in the primary PCa PPI network (**Figure 6B**).Thus the varying pattern of rich club signatures across the hierarchy may possibly relate to the change in popularity of the proteins at different levels of organization, and hence *hub*-proteins preserve their level-dependent influence across the hierarchy (10). Such behaviour in the PPIs networks can be correlated to their weaker resilience and instability at subsystem/modular level which may be critical for certain functional modules due to malfunctions in the key regulator *hub-*proteins.

The Centrality measures are used to assess the importance of the nodes in information processing in the network. Betweenness centrality ***C_B_***, closeness centrality ***C_C_***, Eigenvector centrality ***C_E_***and subgraph centrality ***C_S_*** are the topological properties which can determine efficiency of signal transmission in a network (28,42). The behaviour of these parameters exhibiting power law as a function of degree ***k*** with positive exponents, where the centralities tend to increase with higher degree nodes (**Equation 13)** (**Figure 3**), reveals the increase in efficiency of signal processing with higher degree nodes in PCa network, showing the importance of *hubs* in controlling the flow of information, thereby regulating and stabilizing the network organization. Therefore, *hub*-proteins have a significant influence in regulating the network although they do not control the whole network completely, thereby increasing the risk of being targeted in the network. Hence, the certain *hubs* might be acting as key regulators in PCa and the 19 predicted key regulators might serve as a backbone of the network.

Community detection of the network using Louvain modularity optimization method leads to clustering of the primary PCa network up to the level of motifs (**Figure 2A**). This clustering shows that modularity, ***Q***, of the networks exhibit an increasing pattern with the topological levels with highest average modularity (***Q*** = 0.5527) seen at the first hierarchical level of PCa network and lowest (***Q*** = 0.0013) at level V, that is, at the motif level (43,44). In complex PPI network the modules have biological meanings and gene ontology analyses have revealed enrichment of certain known functions and pathways in the modules (45). The functional homogeneity in the modules of PCa network has been correlated to their mean clustering coefficients as modules with higher mean clustering coefficients have better chance to be associated with specific functions (89,90). Moreover, in disease interactome, the disease modules which are unique modules representing the interaction between disease genes and their neighbourhood, overlaps with the topological modules derived from the network and functional modules associated with functions and are interrelated (91). Primary PCa network is composed of 14 modules deduced from the community detection method with their mean clustering coefficients ***C*(*k*)**∼0.094 − 0.392 (**Table 3**). Among them modules *C*12 and *C*13 which were the largest had the highest mean clustering coefficients ***C*(*k*)** = 0.392 & 0.218, respectively, showing a functional homogeneity in these modules. These modules have been analysed for their functional annotations with DAVID functional annotation tool (40,41) which revealed association with different functions (**Table 3**). Modules *C*12 and *C*13 are represented with ribosomal biosynthesis and transcriptional regulation, respectively. This suggests a bigger role of RPs in PCa which is also evident from the representation of various RPs (*RPL*3,5,6,11,15,19 etc) as key regulators in PCa network. Transcriptional regulation is the most important level of gene regulation which is accomplished mainly through interaction of transcription factors along with their cofactors to the promoter regions of many genes. The tumour suppressor transcription factor (TF) *p53* gene—the most mutated among all PCa—is one of the hub proteins represented in this community. Another important TF, *c-MYC*—an oncogene acting as a regulator of the cell cycle progression and cell division—is also represented in this community. Moreover, reports on regulations of *p53* with the key ribosomal proteins (*RPL*5, *RPL*6, *RPL*11 etc) and *c-MYC* key regulator CUL7 through p53 in several cancers suggest a critical association of transcriptional regulation in PCa.

Since the study of complex hierarchical networks is incomplete without understanding the modularity of subcommunities and the roles played by the nodes in the modules, our study applied the approach to characterize the nodes in PCa network through defining their within-module *Z* score ***Z_i_*** with their participation coefficients ***P*_i_**(36). In the PCa network many *hub* proteins act as *modular kinless hubs* or *connector modular hubs* maintaining the links within the modules as well as connecting other modules at the same level (**Figure 4A,4B,4C**). This shows the importance of the *hub*-proteins in the hierarchical organization of the network exhibiting their involvement in establishing links among the nodes in each module as well as among the modules in the network which are associated with specific functions.

## Conclusions

This paper introduces a new method for finding key regulators in prostate adenocarcinomas using biological networks constructed from high throughput datasets of Prostate cancer patients. The Network theoretical approach used here placed equal emphasis on the hubs, motifs and modules of the network to identify key regulators/regulatory pathways, not restricting only to overrepresented motifs or hubs. It established a relationship between *hubs, modules* and *motifs*. The network used all genes associated with the disease, rather than using manually curated datasets. Highest degree *hubs* (*k* ≥ 65) were identified, out of which 19 were novel key regulators. The network, as evident from fractal nature in topological parameters, was a self-organized network and lacked a central control mechanism. Identification of novel key regulators in prostate cancer, particularly ribosomal proteins add new dimension to the understanding of PCa and its treatment and predicting key disease genes/pathways within network theoretical framework. This method can be used to any networks constructed from patients’ datasets which follow hierarchical topology.

## Acknowledgements

IRM acknowledges Deshbandhu College, University of Delhi for study leave to pursue doctoral research. MZM was financially supported by the Department of Health and Research, Ministry of Health and Family Welfare, Government of India under young scientist FTS No. 3146887. SA acknowledges DBT, Ministry of Science and Technology, Government of India for Bioinformatics BIF grant under the Biotechnology Information System Network (BTISNET), sanction number BT/BI/25/062/2012(BIF). RKBS acknowledges financial support by UPE-II under sanction no. 101, India.

## Author Contributions

RKBS, SA, IRM and MZM conceived the model and conducted numerical experiments. IRM and MZM prepared figures of the numerical results. IRM, MZM, OK, SA and RKBS analysed and interpreted the simulation results and wrote the manuscript.

---

Supplementary Table 1. Top 103 *hubs*.

